# Early hominin arrival in Southeast Asia triggered the evolution of major human malaria vectors

**DOI:** 10.1101/2024.09.28.615606

**Authors:** Upasana Shyamsunder Singh, Ralph E. Harbach, Jeffery Hii, Moh Seng Chang, Pradya Somboon, Anil Prakash, Devojit Sarma, Ben Broomfield, Katy Morgan, Sandra Albert, Aparup Das, Yvonne-Marie Linton, Jane M. Carlton, Catherine Walton

**Affiliations:** Department of Earth and Environmental Sciences, School of Natural Sciences, University of Manchester, Manchester, United Kingdom; Department of Biological Sciences, Vanderbilt University, Nashville, USA; Department of Science, Natural History Museum, Cromwell Road, London, UK; College of Public Health, Medical and Veterinary Sciences, James Cook University, North Queensland, Australia; Department of Community Medicine & Public Health, University Malaysia Sarawak, Malaysia; Center of Insect Vector Study, Department of Parasitology, Faculty of Medicine, Chiang Mai University, Chiang Mai, Thailand; ICMR-Regional Medical Research Centre, Dibrugarh, Assam, India; ICMR-National Institute for Research in Environmental Health, Bhopal, India; Indian Institute of Public Health Shillong, Shillong, Meghalaya, India; ICMR-National Institute of Research in Tribal Health, Jabalpur, Madhya Pradesh, India; Walter Reed Biosystematics Unit, Smithsonian Museum Support Center, Suitland, MD, USA; Department of Entomology, Smithsonian Institution – National Museum of Natural History, Washington, DC, USA; One Health Branch, Walter Reed Army Institute of Research, Silver Spring, MD, USA; Johns Hopkins Malaria Research Institute, Bloomberg School of Public Health, Baltimore, MD, USA

## Abstract

Understanding the evolution of anthropophily, the preference of mosquitoes to feed on humans, offers insights into current and future human disease transmission. Some species of the Leucosphyrus Group of *Anopheles* mosquitoes in Southeast Asia are highly anthropophilic and efficient vectors of human malaria parasites, while others primarily feed on non-human primates and transmit non-human primate malaria parasites. Through phylogenomic analysis of 11 out of 20 recognized species, we studied the biogeography and evolutionary history of anthropophily in this group. Molecular dating and ancestral state reconstruction revealed that anthropophily evolved during the late Pliocene/early Pleistocene in Sundaland, likely in response to early hominins. This finding provides independent non-archaeological evidence supporting the limited fossil record of early hominin colonization in Southeast Asia around 1.8 million years ago.

## Introduction

Mosquito-borne diseases present a significant burden on human health, with malaria alone causing an estimated 249 million cases and 608,000 deaths worldwide in 2022 (*1*). The propensity of mosquitoes of a particular species to feed on humans (anthropophily) is the primary factor influencing their potential to spread pathogens that cause disease (*2–7*). Although mosquitoes can be opportunistic in their host selection (*e.g., 8, 9*), many species display varying degrees of host specificity (*10, 11*). Understanding the evolutionary origins of anthropophily and the circumstances that triggered its development can provide critical insights into mitigating the impacts of novel diseases due to mosquito-borne pathogens.

The *Anopheles leucosphyrus* group (hereafter, Leucosphyrus Group) comprises 20 recognized mosquito species in Southeast Asia (SE Asia) (*12–15*). These species exhibit intrinsic differences in host preference, as demonstrated by host attraction experiments, blood-meal analysis, and variation in transmission of human and non-human primate (NHP) malarias (table S1) (*13, 16–19*). Notably, several species are highly anthropophilic and extremely efficient vectors of human malaria parasites. These include *An. dirus, An. baimaii,* and *An. scanloni* of the Dirus Complex found in mainland SE Asia, and *An. balabacensis* of the Leucosphyrus Complex from Borneo (Sabah and Kalimantan) (*17, 20–23*) (table S1). Conversely, species such as *An. macarthuri*, *An. pujutensis*, and *An. hackeri* blood-feed only in the forest canopy on NHP, including monkeys, gibbons, and orangutans, transmitting NHP malaria parasites (Fig. 1, table S1) (*24–27*). *Anopheles nemophilous* (of the Dirus complex), *An. latens*, and *An. introlatus* (of the Leucosphyrus Complex) are less host-specific, feeding on both NHPs in the canopy and humans on the ground, apparently driven by host availability (table S1). As host choice experiments most often compared humans on the ground, to monkeys in the canopy, it is not possible to separate a tendency to seek hosts on the ground rather than in the canopy as a distinct trait (*23, 28, 29*) (table S1).

**Fig. 1.**
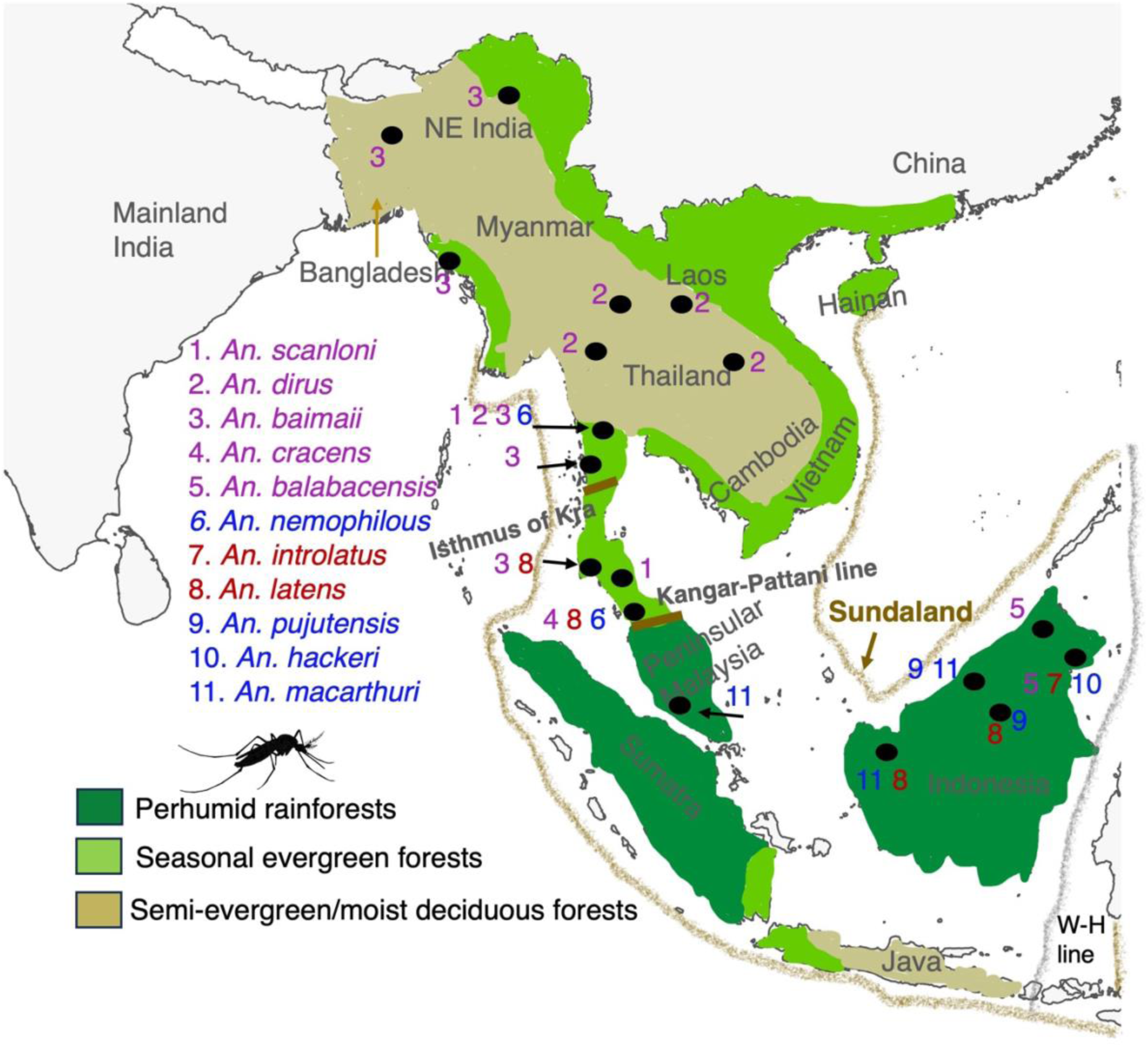
Map representing the distributions of specimens collected in Southeast Asia. Shading indicates the present-day distributions of forest types in mainland and insular Southeast Asia, adapted from Morley (*49*). Black dots on the map represent collection sites. The number adjacent to the dots represents individual species collected from that site according to the species list on the left. The color of the numbers and species names indicate distinct blood-feeding behaviors; blue–NHP feeding, red–mixed-feeding, purple–human feeding, derived from published literature (listed in table S1). The brown outer line represents the outline of the exposed Sunda Shelf at the Last Glacial Maxima (currently 120 m below sea level) (*52*). The two short brown lines represent biogeographic barriers: the Kangar-Pattani line in the south and the Isthmus of Kra in the north. The Isthmus of Kra and the W-H (Wallace-Huxley) line (grey line) in the east mark the boundaries of Sundaland.

The establishment of anthropophily in multiple species of the Leucosphyrus Group could be attributed to the trait evolving independently multiple times following the arrival of anatomically modern humans in SE Asia 76,000–63,000 years ago (*30, 31*). Alternatively, anthropophily may have evolved once in an ancestral species, possibly in response to the colonization of SE Asia by early hominins. Conservative estimates place *Homo erectus* in China at least 1.6–1.7 million years ago (Mya), and possibly as long ago as 2.4 Mya (*32*). However, the timeline of hominin colonization southwards into SE Asia remains contentious. Recent reports suggest that hominins may have arrived in Java between 1.3 (*33*) and 1.8 Mya (*34*). Increased aridity during the Late Pliocene and Early Pleistocene, particularly during periodic glacial periods, is considered to have resulted in the formation of a north-to-south corridor of seasonal forests and grasslands (*35*), that facilitated early hominin migration through SE Asia into Java (*36*). We used phylogenomics and analyses of trait evolution in the Leucosphyrus Group to characterize the evolutionary history of these mosquitoes in relation to historical environmental changes and host preference. Our findings offer independent, non-archaeological evidence for the timing and location of the early hominin colonization of SE Asia, providing new perspectives on the co-evolution of mosquitoes and their hosts.

### Genome-scale phylogenies reveal reticulate evolution in the Leucosphyrus Group

To elucidate the evolutionary history of the Leucosphyrus Group, we sequenced 38 individual mosquitoes of 11 species, supplemented with the publicly available genomes of *An. dirus* (*37*) and *An. cracens* (*38*). Many of these species are particularly challenging to collect, for example, involving sampling larvae from animal wallows deep in the forest and from remote locations. Specimens of the 11 species studied here were accumulated over several years (from 1992-2020) and include all species of the Leucosphyrus Group from Sundaland and Indochina except, for logistical reasons, those restricted to Sumatra and the Philippines. These 11 species include members of all three Subgroups (Leucosphyrus, Riparis and Hackeri) and represent all three blood-feeding behaviors (human, NHP and mixed human-NHP) (Fig. 1, table S2). Orthology inference using *An. dirus* and an outgroup species, *An. farauti*, identified 2,657 high-confidence nuclear single-copy orthologous genes (nSCOs) across 40 genomes. Phylogenetic reconstructions were performed using coalescent-based summary analyses with ASTRAL (*39*) and maximum-likelihood (IQ-TREE), (*40*) analyses on 2,657 nuclear and 13 mitochondrial protein-coding DNA sequences (Fig. 2).

**Fig. 2.**
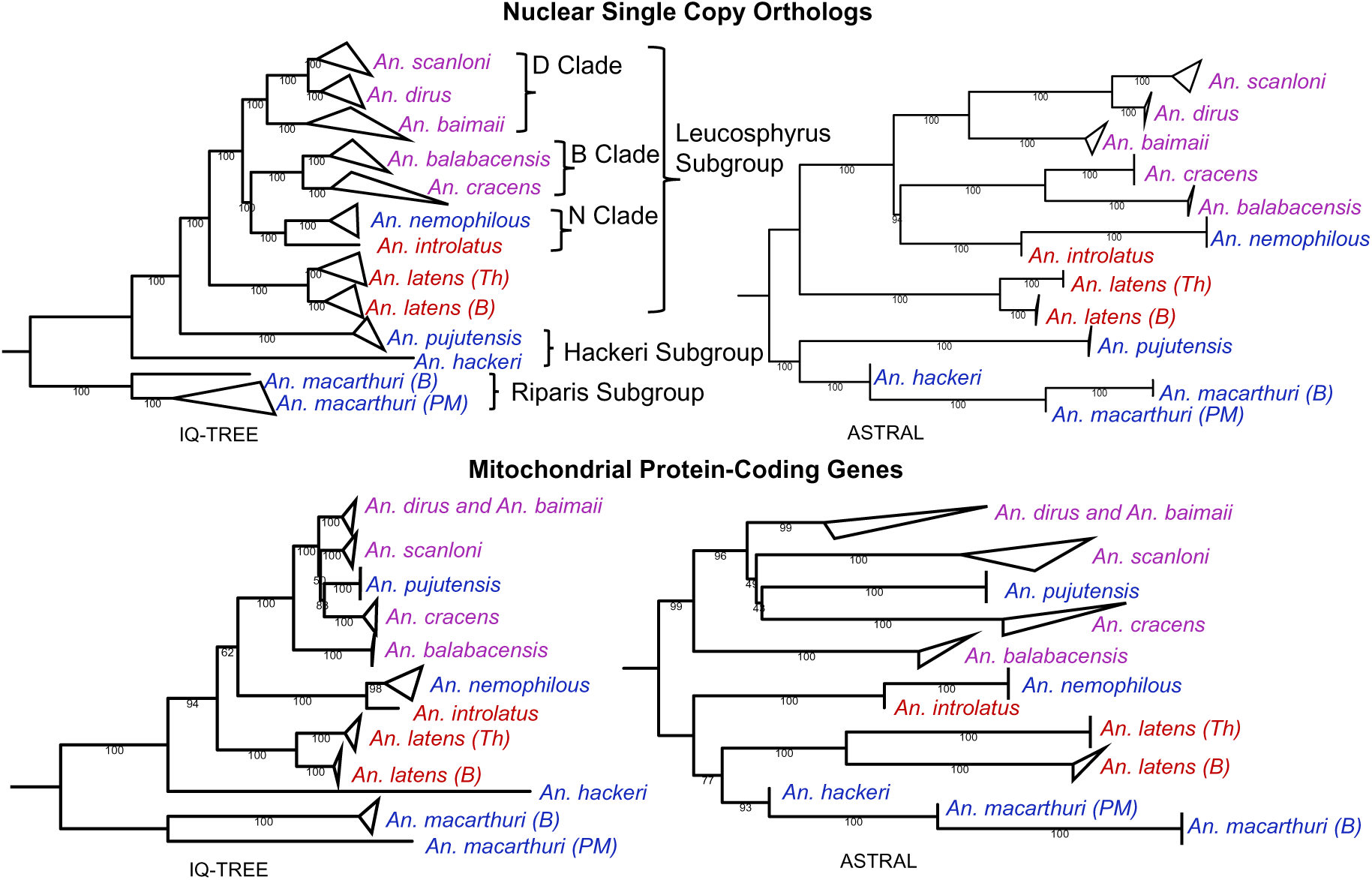
Genome-scale phylogenies of the Leucosphyrus Group. Trees were constructed using concatenation-based (IQ-TREE) (right) and ASTRAL (left) approach. The nuclear trees (top) were constructed using 2,657 single copy orthologs. The mitochondrial trees (bottom) were constructed using 13 protein-coding genes. Numbers on the branches denote bootstrap values. Species names are color-coded according to feeding preferences as in Figure 1 and table S1. Based on the morphological classification by Sallum et al. (*17*), the Leucosphyrus Group comprises the Riparis, Hackeri and Leucosphyrus Subgroups. The Leucosphyrus Subgroup is further divided into the Dirus Complex (*An. baimaii*, *An. dirus*, *An. scanloni*, *An. cracens* and *An. nemophilous*) and the Leucosphyrus Complex (*An. balabacensis, An. latens* and *An. introlatus*) (*17*). The morphological classification and phylogenetic classification do not agree therefore, based on the nuclear phylogeny, we classify *An. dirus*, *An. baimaii* and *An. scanloni* as the Dirus Clade (D Clade), *An. balabacensis* and *An. introlatus* as the *Balabacensis* Clade (B Clade) and *An. cracens* and *An. nemophilous* as the Nemophilous Clade (N Clade). Different population of *An. latens* and *An. macarthuri* are denoted by: Th–Thailand, B–Borneo and PM– Peninsular Malaysia.

Inconsistencies between the resultant nuclear and mitochondrial phylogenies (Fig. 2) indicated mitochondrial introgression. *Anopheles dirus* and *An. baimaii*, though distinct in the nuclear phylogenies, are indistinguishable in mitochondrial phylogenies, confirming recent mitochondrial introgression (*41*). Based on the placement of *An. pujutensis* (Hackeri Subgroup) within the Dirus Complex in the mitochondrial phylogenies, and its placement with other NHP-feeding species in the nuclear phylogenies, we infer older mitochondrial introgression from a member of the Dirus Complex into *An. pujutensis* (Fig. 2). *Anopheles cracens* is also differently placed between the mitochondrial and nuclear trees (Fig. 2). Due to this mitochondrial introgression, we focused subsequent analyses on the nuclear data.

The tree topologies for each genomic region (nuclear or mitochondrial) were consistent across phylogenetic methods, except for the placement of a midpoint root among *An. macarthuri*, *An. hackeri*, and *An. pujutensis*, indicating variation in branch length estimation between ASTRAL and IQ-TREE methods. Discordance between coalescent-based (ASTRAL) and concatenation-based (IQ-TREE) methods is commonly observed and can be attributed to multiple causes, including differing evolutionary histories among loci, rate variation among lineages, or model misspecification (*42, 43*). To investigate these discrepancies, we used PhyloNet (*44*) to construct phylonetworks. As the number of reticulations increased from one to three, the likelihood of the phylonetworks increased (fig. S1). Although more than three reticulations might be likely, further exploration was limited by computational power. All networks with 1– 3 reticulations indicated substantial introgression of nuclear loci between lineages leading to *An. pujutensis, An. hackeri*, and *An. macarthuri*, and some introgression between these lineages and more recently derived species (fig. S1). These introgressions may significantly contribute to discrepancies in the placement of the midpoint root.

### Evolutionary timeline and geographical origin of mosquito feeding preference

Overall, the nSCO phylogenies challenge the current morphology-based taxonomic classification (*17*). While there is support for the monophyly of the Leucosphyrus Subgroup, the phylogenies do not support the monophyly of the Dirus and Leucosphyrus Complexes within this Subgroup (Fig. 2). Despite uncertainty in the order of basal branching due to introgression, the high topological concordance of the nuclear phylogenies provides a robust phylogenetic framework to study the evolution of host preference for species of the Leucosphyrus Group. Further, to make the analysis computationally tractable and minimize lineage rate variation among species, we identified 25 clock-like genes (*45*) for reconstructing a chronogram in BEAST (Bayesian Evolutionary Analysis Sampling Trees) (*46*) (Fig. 3). This phylogeny was used to estimate divergence time (Fig. 3), and to reconstruct the ancestral states for host preference (Fig. 4a) and biogeographical distribution (Fig. 4b) using Reconstruction of Ancestral States (RASP) (*47*).

**Fig. 3.**
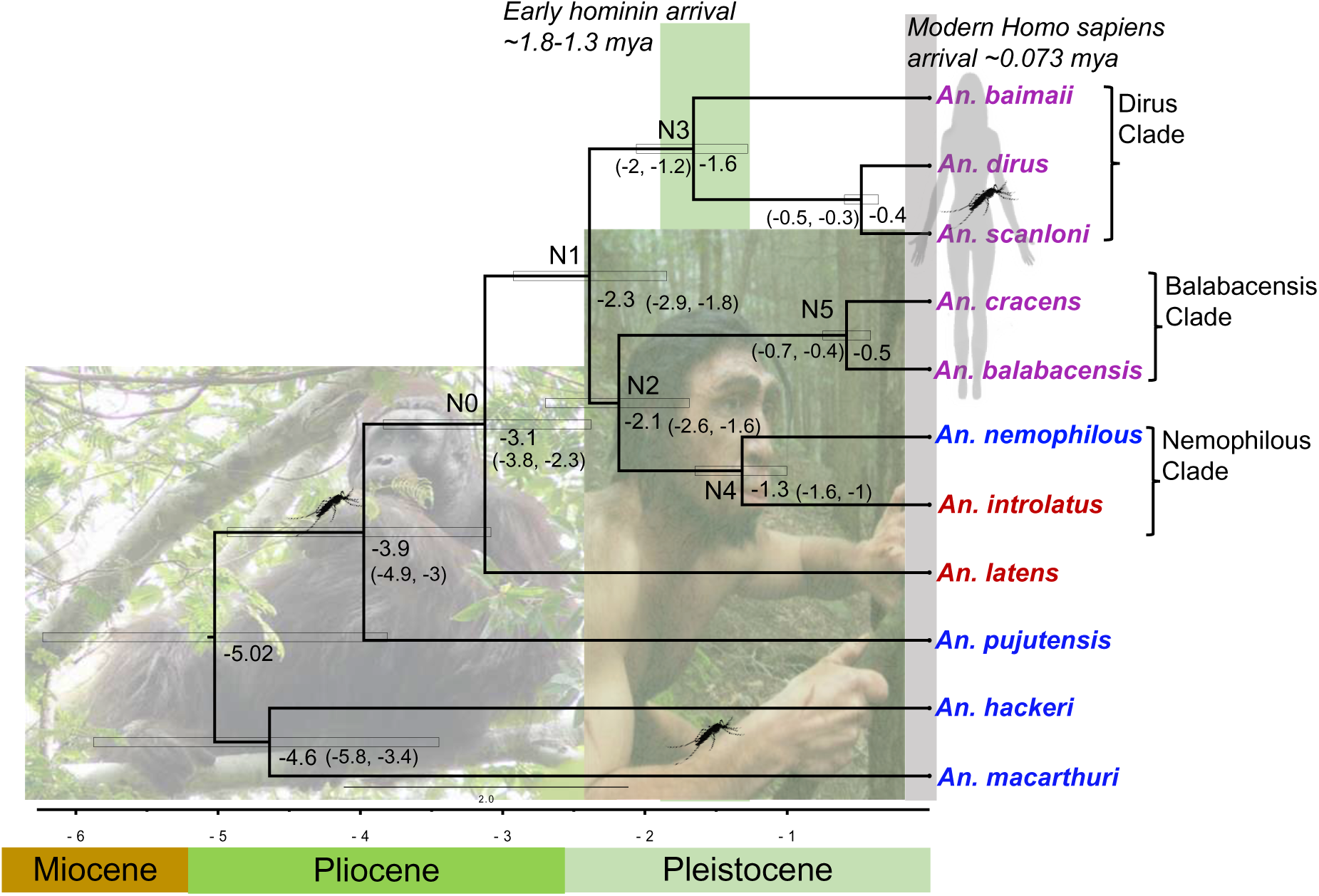
Chronogram of the Leucosphyrus Group using a Bayesian approach. The divergence dating analysis was carried out using 25 clock-like genes and the coalescent species tree estimation method. Colors of the named species indicate host feeding preferences (see Fig. 1 and table S1). The estimated divergence times are indicated by the nodes with the 95% highest posterior densities denoted by blue bars and the numbers in parentheses. The green vertical shading represents the time interval proposed for the arrival of early hominins in Southeast Asia based on fossil data between 1.3 (*33*) and 1.8 million years ago (*34*). The vertical grey line represents the time of arrival of anatomically modern humans (*30*).

**Fig. 4.**
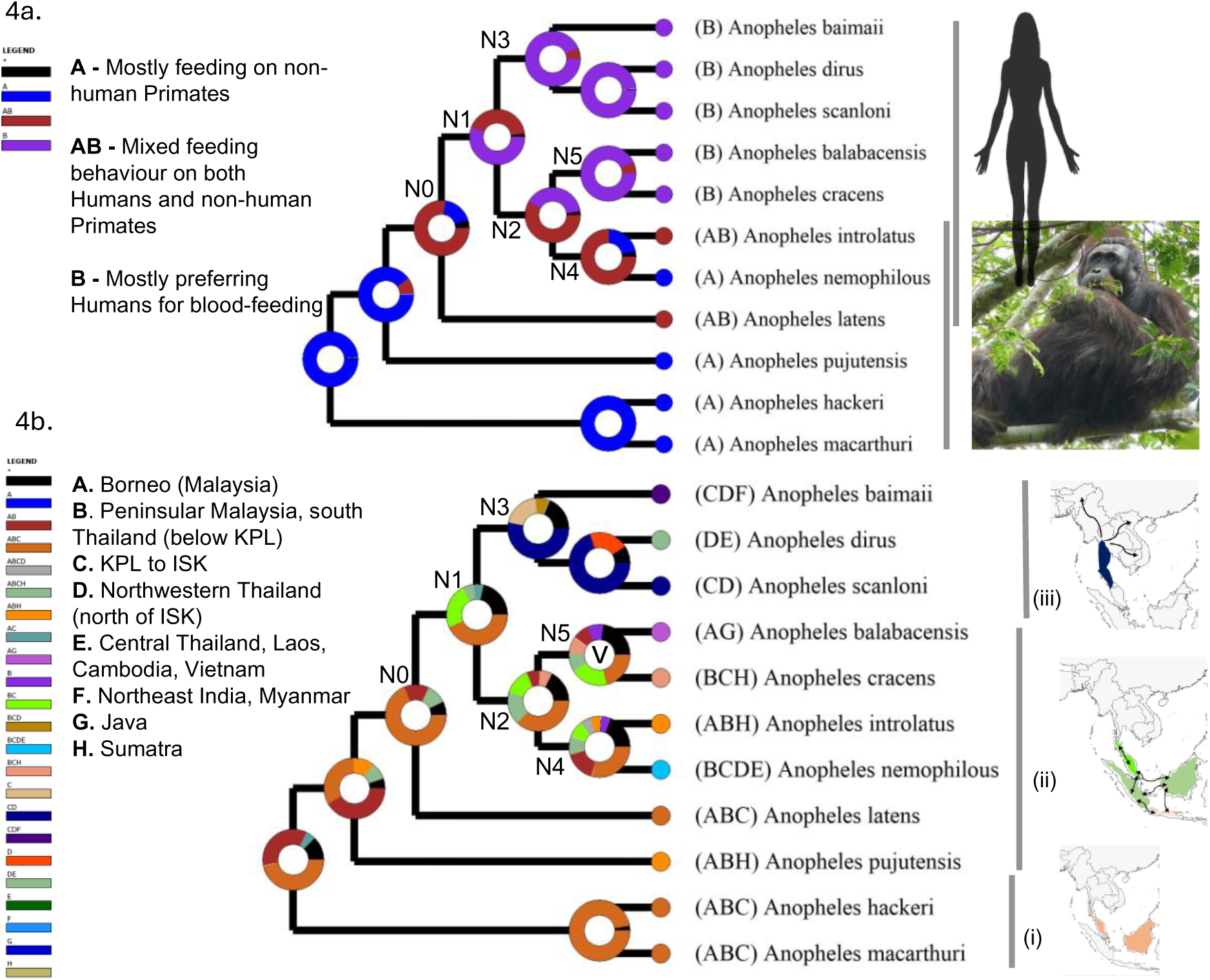
Reconstruction of ancestral states for blood-feeding behavior and biogeography. **(4a)** Trait evolution analysis of host preference for blood-feeding. The node N0 represents a switch from feeding only on NHPs in the canopy to also feeding on humans on the ground. Node N1 represents the earliest inferred switch to a preference for feeding on humans. Node N3 represents the ancestor of the Dirus Clade; N4–ancestor of the Nemophilous Clade, and N5–ancestor of the Balabacensis Clade. **(4b)** Reconstruction of the ancestral states for biogeography. This analysis shows dispersal at every node except for node N5 where “V” inside the pie denotes a vicariance event. The schematic maps on the right (**i–iii**) show dispersal within the Leucosphyrus Group for the corresponding part of the phylogeny, where (**i**) the orange color shows that peninsular Malaysia and Borneo is the ancestral range of monkey-feeding groups, (**ii**) green shows subsequent dispersal between Peninsula Malaysia, Sumatra, Borneo, and Java followed by (**iii**) shown in blue, dispersal of the Dirus Clade to and across the Indochinese region. Black arrows indicate direction of dispersal.

*Ancestors of the Leucosphyrus Group were feeding on non-human primates in Sundaland* Extensive introgression among *An. macarthuri*, *An. hackeri*, and *An. pujutensis* complicates the precise estimation of divergence times however, we infer that these NHP-feeding species are basal and diverged during the early Pliocene (5.3–3.6 Mya) (Fig. 3). Ancestral trait reconstruction for host preferences (Fig. 4a) corroborates the expectation that NHP-feeding is the ancestral state. Additionally, biogeographical reconstruction (Fig. 4b) indicates these three species originated in Sundaland (Fig. 1), encompassing present-day Borneo, peninsular Malaysia, peninsular Thailand below the Isthmus of Kra, Sumatra, and the currently submerged Sunda Shelf. During this period, the region was covered by extensive permanently humid (perhumid) rainforests (*48, 49*), providing ample opportunity for specialization in feeding on NHPs in the forest canopy.

### Speciation associated with Plio-Pleistocene environmental change

The Pliocene and early Pleistocene were characterized by increasingly cooler and drier global climates (*50, 51*). It is during this period, characterized by extensive environmental change, that the Leucosphyrus Subgroup emerged, with the ancestral species *An. latens* diverging at node N0, around 3.1 Mya [95% CI: 3.8–2.3 Mya] (Fig. 3). Subsequently, four divergence events (nodes N1–N4, Fig. 3) occurred in the early to mid-Pleistocene (2.3–1.3 Mya) within Sundaland (Fig. 4b), with divergence at N1 giving rise to the Dirus Clade (comprising *An. dirus, An. baimaii and An. scanloni*) and divergence at N2 giving rise to the Nemophilous (*An. nemophilous* and *An. introlatus*) and Balabacensis (*An. balabacensis* and *An. cracens*) Clades (Fig. 2, 3). It has long been viewed that eustatic sea-level changes were the major driver of diversification during the Pleistocene, with repeated splitting and reformation of the Sundaland landmass (*48*, *49*, *52*). This view has changed with the recognition of subsidence of the Sunda Shelf during the Pleistocene, which means that it must have been exposed as a single landmass continuously until 400,000 years ago (*53*). It is only after 400,000 years that Java, Sumatra, Borneo, and Indochina–Peninsular Malaysia were separated by elevated sea levels during interglacial periods (Fig. 1). Consequently, throughout the late Pliocene and most of the Pleistocene the balance of evidence favors a seasonal corridor that extended from Indochina southwards through central Sundaland, featuring more open and seasonal forests and potentially including grasslands particularly during drier glacial periods (*35, 54–56*). Therefore, we propose that the apparent burst of speciation occurring at nodes N0, N1–N4 within Sundaland at this time involved adaptation to novel forest types, hosts associated with these new habitats, or vicariance involving repeated fragmentation of perhumid rainforests. Since bi-directional dispersal was detected (Fig. 4b) between present-day peninsular Malaysia, Sumatra, Borneo, and Java, it indicates that there may have been limited periods of forest connectivity across Sundaland (*49*). The recent divergence of *An. cracens* and *An. balabacensis*, which are restricted to peninsular Malaysia and Borneo, respectively, at node N5 around 0.5 Mya [95% CI: 0.7–0.4 Mya] (Figs. 3, 4b), is attributed to vicariance by the RASP analysis, likely due to the formation of separate landmasses during interglacial periods.

### Crossing the barrier of the Isthmus of Kra and origin of the Dirus Clade

The Isthmus of Kra (ISK) marks a significant biogeographical boundary for many forest species, including birds (*57*), due to the seasonally dry climates in lowlands further north of ISK, in Thailand (*49*). While *An. nemophilous* is found both north and south of the ISK barrier, its distribution is confined to seasonal evergreen forests south of the ISK, and above the Kangar-Pattani line (K-PL) and patches found on the Thai-Cambodia border (Fig. 1) (*20*). Speciation of the ancestor of the Dirus Clade likely involved adaptation to the seasonal forests north of ISK that enabled it to cross this biogeographical boundary. This ancestral species migrated northward, where it subsequently gave rise to at least three species (*An. dirus*, *An. baimaii*, and *An. scanloni*), with *An. dirus* and *An. baimaii* dispersing extensively eastwards and westwards, respectively (Fig. 4b, 4c(iii)), becoming major vectors of human malaria parasites across much of Indochina (*58*).

### Two-stage transition to feeding on humans in Sundaland

The shift away from strict canopy feeding on NHPs is inferred to have begun in the Late Pliocene (Fig. 3), with the emergence of the Leucosphyrus Subgroup (node N0, Figs. 3, 4a). Unlike its canopy-feeding ancestors, the basal species of this Subgroup, *An. latens*, readily feeds both on humans on the ground and NHPs in the canopy (*24, 26, 59–61*) (table S1). The abundance of ground-dwelling mammals likely increased with the transition from perhumid to seasonal and open forest types (*49, 62*). Consequently, the evolutionary change in host-seeking behavior of *An. latens* likely represented an adaptive evolutionary innovation, involving a willingness to seek hosts on the ground.

### Evolution of vectors of malaria parasites in response to early hominin colonization of Southeast Asia

Mosquitoes employ multiple senses to track their hosts, but evolutionary changes in olfactory genes, particularly those involving the fine tuning of olfactory receptors through modification of their expression and specificity, are crucial for developing a preference for human body odor (*63*). The large number of odorants and olfactory genes involved in host specificity indicates that multiple genetic changes are required for the evolution of anthropophily. Therefore, it is improbable that there were five independent switches to anthropophily in the human-preferring species of the Dirus and Balabacensis Clades, which diverged around 1.3–0.5 Mya (Fig. 3).

Based on our molecular clock dating, we reject the hypothesis that anthropophily in the Leucosphyrus Subgroup evolved in response to the arrival of anatomically modern humans in SE Asia around 76,000–63,000 years ago (*30, 31*). Instead, Figure 3 indicates that anthropophily emerged by the time of node N1 [95% CI: 2.9–1.8 Mya] and was subsequently lost in the lineage leading to *An. nemophilous* and *An. introlatus*. This confidence interval only slightly overlaps with the earliest proposed date for the arrival of early hominins at 1.8 Mya (*34*), suggesting that anthropophily evolved in response to the arrival of early hominins in Sundaland. Genetic introgression can occur throughout an extensive speciation process (*64*), especially if associated with adaptation (*65*). An alternative hypothesis for the shared anthropophily of the paraphyletic Balabacensis and Dirus Clades (Fig. 3, nodes N3 and N5) is therefore that genetic changes associated with anthropophily introgressed between the lineages leading to these Clades. This is most likely to have occurred after divergence at node N1 (Figs. 3, 4) when all the lineages were in Sundaland, and prior to the divergence of the Dirus Clade (node N3, 1.6 Mya [95% CI: 2.0–1.2 Mya]) further north in Indochina.

Based on dating the emergence of anthropophily in other mosquito species, *e.g*., the domestic form of *Aedes aegypti* (*66*); the molestus ecotype of *Culex pipiens* (*7*), and the major African vectors *An. gambiae* (*67*), the abundance of a novel host source appears to be a key requisite for triggering host-switch. Our findings suggests that anthropophily in the Leucosphyrus Group emerged in Sundaland in the early Pleistocene ∼1.8 Mya, indicating that early hominins were not only present in this region at this time but must have been in substantial numbers to drive adaption to human host preference. This supports the hypothesis of Husson et al. (*34*) that early hominins were present and abundant in Sundaland ∼1.8 Mya, prior to their dispersal via land bridges to Java.

## Methods and Supplementary Materials

### Materials and Methods

#### Fieldwork and Sampling/Taxon Sampling

Mosquito specimens (n = 38) were used to represent 11 of the 20 species in the Leucosphyrus Group, including species belonging to each Subgroup and representing all blood-feeding behaviours (table S2). Except for colony material of *An. cracens*, all the specimens were obtained as either larvae or adults from field collections between 1995 and 2020 (table S2). Most adult mosquitoes were collected using human landing catch, while *An. pujutensis*, *An. macarthuri*, *An. hackeri*, *An. introlatus* and one *An. balabacensis* and one *An. baimaii* were collected as larvae (table S2). The larvae were reared to adults for morphological identification to species or as belonging to the *An. dirus* complex using keys by Reid (*13*) and E L Peyton (unpublished).

### DNA extraction, library preparation and sequencing

DNA extractions of whole mosquitoes were carried out following a phenol-chloroform protocol. Total genomic DNA extraction for a few mosquito specimens was performed with DNeasy Blood and Tissue Kits (Qiagen®), digested overnight with proteinase K following the manufacturer’s protocol, and eluted with elution buffer to either 50 or 100 µl. Extracted genomic DNA was quantified with a Qubit 4.0 fluorometer (Invitrogen, Carlsbad, CA, USA) using the manufacturer’s protocol and 20 µl of DNA (10-100ng/µl) per sample was sequenced at the Earlham Institute (Norwich, UK). Low Input, Transposase Enabled (LITE) libraries were constructed and sequenced on two lanes of the NovaSeq 6000 SP flow cell with 150bp paired-end reads.

### Quality control of raw reads

With a sequencing depth of ∼30X, we were able to generate 3 million paired-end DNA sequence reads and roughly 10 GB of data per sample. To minimise sequencing errors several quality control steps were taken to filter out low-quality sequencing reads. Raw FASTQ reads were filtered using the software TrimGalore (*68*) by trimming adapters and low-quality bases (Phred quality ≥30) and cleaned sequences were reanalysed using FastQC (*69*) and MultiQC (Babraham Bioinformatics).

### Single-copy ortholog identification and filtering

The assembled and annotated proteomes of the reference species *An. dirus s.s.* (ENA accession-GCA_000349145.11.1) from Thailand and an outgroup species *An. farauti* (ENA accession-GCA_000473445.2) from Papua New Guinea were used to define groups of orthologous sequences using OrthoFinder2 (*70*). Only single-copy orthologs (SCOs) that were present in all individuals and had a minimum length of 300-bp were chosen for downstream analysis, which resulted in 5,867 nuclear SCO protein-coding genes.

### Assembly, alignment and filtering of nuclear SCOs

We used aTRAM 2.0 (*71*), an iterative assembler that executes reference-guided local *de novo* assemblies, to assemble 5,867 SCOs from 38 genomes we sequenced. To do this, we used trimmed sequence reads that were first converted to a Blast database using the atram_preprocessor.py command of aTRAM v2.3.4. The amino acid sequences of 5,867 genes were used in tblastn along with the SPAdes assembler (*72*), with five iterations, to create the aTRAM assemblies (atram.py script). The exon sequences from aTRAM assemblies were stitched together in the correct frame using the find_orthologs.py wrapper script (*73*). Two publicly availsble genomes, one for *An. dirus s.s.* (GCA 000349145.1, Thailand) (*37*) and one for *An. cracens* (GCA 002091845.1, peninsular Malaysia) (*38*), were used in addition to the 38 novel genomes for phylogenetic analysis. From the 5,867 genes that were assembled by aTRAM we selected 2,928 that were present in all the samples for phylogenetic analysis. Each gene was aligned using the – auto flag in MAFFT (*74*) and individual gene alignments were trimmed using the-automated1 flag in trimAl v. 1.4.1 (*75*). Gene alignment of length >500bp was chosen for phylogenetic analysis. This filtering resulted in 2,657 SCOs.

### Mitochondrial genome assembly and alignment

The mitochondrial genome was assembled using MITObim v1.9.1 program (*76*). First, trimmed paired-end reads were interleaved using NGmerge v0.3 (*77*). The complete mitochondrial genome from *An. dirus s.s*. (GenBank accession NC_036263) was used as a reference seed, and the assemblies were created with 30 iterations and --quick option using the Mitobim.pl script. The FASTA reads retrieved in the final iteration were annotated using default parameters in the MITOS web server (*78*) (http://mitos.bioinf.uni-leipzig.de/). The GFF files generated by MITOS were imported to Geneious Prime v11.0.4. and 13 mitochondrial protein coding genes (PCGs) were extracted from mitogenomes. Two published mitogenomes of *An. dirus s.s.* NC_036263 (Hainan, China) and *An. cracens*, NC_020768 (Thailand), were also used along with 38 samples. Geneious v11.0.4’s MUSCLE aligner was used to align PCGs from 40 individuals, and trimming was performed so that all genes were the same length.

### Sample validation

To validate the species identity and identify any potential contamination in our assembled sequences, we used the NCBI BLAST web interface to compare our ITS2 sequences assembled in aTRAM against the GenBank database. We also verified the *COI* sequences assembled using MITObim against BOLD and GenBank databases. Specifically, rDNA *ITS2* was used to distinguish *An. dirus* from *An. baimaii* as they cannot be separated using mtDNA *COI* (*41*). Morphological identification for *An. hackeri*, *An. pujutensis*, *An. macarthuri*, *An. balabacensis* was performed in the field by an expert taxonomist (Ralph E. Harbach).

### Phylogenetic analysis

Two datasets were used to reconstruct the phylogeny of eleven species in the Leucosphyrus Group: mitochondrial PCGs and nuclear SCOs of 40 individuals. Both concatenation and coalescent-based methods were used. We first concatenated all 13 PCGs in Geneious to make a bigger alignment file consisting of 9,900 bp. The nuclear dataset was made by concatenating 2,657 SCOs using the script concatenate.rb (*73*) to make a supermatrix of 4,929,412 bp. Maximum likelihood phylogenetic reconstruction using the concatenated dataset was performed in IQ-TREE v2.1.2. (*40*). We ran MODELFINDER (*79*) in IQ-TREE with the –m MFP option to find the best model. The best-fit model: TIM2+F+I+G4F was chosen for mitochondrial PCGs and GTR+F+R10 was chosen for the nuclear dataset according to Bayesian Information Criterion. Incorporating these models in the respective datasets, 100 ultrafast bootstraps were performed in IQ-TREE v2.1.2. For the coalescent-based tree reconstruction, we first generated gene trees for each gene using 100 rapid bootstrap replicates in RAxML v8.2 (*80*) using a GTRGAMMA model. The gene trees for both the mitochondrial and nuclear datasets were then summarised using ASTRAL v5.7.8. Both the trees were visualised using iTOL (*81*).

### Divergence time estimation

The Leucosphyrus Group divergence time was estimated using the Bayesian phylogenetic method in BEAST v2.6.7 (*46*). However, due to the computational demands of the Bayesian approach, we were unable to make use of the entire nuclear dataset. We used gene selection to create a reduced dataset that could be processed with the available computing power and time. In a study by Jarvis et al. (*82*), “clock-like” genes were found to evolve at a steady rate and reduce errors caused by model misspecification (*45*). Therefore, we utilised SortaDate to find genes based on root-to-tip variation, bipartition, and tree length (in order) (*45*). We selected 25 clock-like nSCOs nucleotide alignments to create a dataset for dating analysis. Chronograms for nSCOs were generated using StarBeast3 (*83*) and standard BEAST templates (*46*). As no mutation rate is available for *Anopheles*, we estimated divergence times using the mutation rate estimated from spontaneous mutations in whole genomes of *Drosophila—* 2.8×10^-9^ mutations per site per generation with 11 generations per year (*84*). To account for only protein coding regions being used here, this rate was scaled by 0.524 based on the relative mutation rate of autosomal coding regions in *Anopheles* (*67*) to yield a rate 1.47×10^−9^ mutations per site per generation. The optimal substitution model, yielding GTR + 4 gamma count categories, was inferred using bModelTest (*85*) in BEAST v2.6.7. Additionally, .xml configuration files with a strict clock were generated using BEAUti v2.6.7. Two independent MCMC runs were carried out, each with a chain length of 100,000,000 and logging every 50,000 generations, until the effective sample size reached above 200. To determine whether each run converged, we used Tracer v1.7.2 to visually evaluate traces and effective sample sizes for posteriors, and likelihoods. We used the TreeAnnotator application in BEAST v2.6.7 (*46*) to generate a maximum clade credibility tree. Trees were visualised in FigTree v1.4.4 (*86*).

### Ancestral state reconstruction

To infer historical biogeography of the Leucosphyrus Group including the relative roles of vicariance and dispersal, the ancestral states were reconstructed on a phylogenetic tree with the RASP (Reconstruct Ancestral State in Phylogenies) v4 software (*87*) using Bayesian Binary MCMC (BBM) analysis. The Bayesian tree generated for divergence dating estimation was used to make a consensus tree. The geographical distributions of the Leucosphyrus Group species (table S3) were based on the literature (*17*) (table S1) with each species assigned to one or more of the following eight geographical areas that capture biogeographical and landmass transitions: (A) Borneo, (B) Peninsular Malaysia + south Thailand below the Kangar-Pattani Line (K-PL), (C) from K-PL northwards to the Isthmus of Kra (ISK), (D) from ISK to northwestern Thailand along its border with Myanmar, (E) remaining area of Thailand + Laos + Vietnam + Cambodia, (F) Myanmar + northeast India, (G) Java, and (H) Sumatra. RASP analysis was also performed to track evolution of the trait for host preference in the Leucosphyrus Group. For this, three categories for trait state were used: (A) non-human primate feeders, (B) mostly anthropophilic, with a third category (AB) indicating feeding on both humans and NHPs without a strong preference, based on evidence from the literature outlined in table S1. For both historical biogeography inference and host preference trait analysis, the MCMC chains of the BBM analysis were run for one million generations, with a sampling frequency of every 100 generations and a 10% burn-in. A fixed JC + G (Jukes-Cantor + Gamma) model was used for the BBM analysis.

### Introgression analysis

Using PhyloNet v3.8.0 (*44*), we reconstructed phylogenetic networks that allow for reticulation between branches. The input comprised of gene trees with one representative individual per species. PhyloNet can infer species networks based on topology alone however, simulations conducted by Yu et al. (2014) have demonstrated that the accuracy of the inferred species networks can be enhanced by incorporating branch lengths. While using branch lengths, PhyloNet also requires the tree to be ultrametric therefore, input trees were generated in BEAST2 (*46*), which produces ultrametric trees, with branch lengths. The gene trees were estimated for 2,657 SCO using the R package babette (*89*) specifying the HKY model of sequence evolution and a chain length of 500,000 iterations. The R script convert_to_newick.r (*73*) was used to convert the Maximum clade credibility trees produced by BEAST2 for each SCO alignment into Newick format, to combine them into a single input file for PhyloNet. Finally, network searches were performed using “InferNetwork_ML” using branch lengths of gene tree (-bl), allowing 0–3 reticulations and optimising branch lengths and inheritance probabilities (-o). Five iterations were carried out and the log probability of the best inferred network was used to compare log probabilities for 1–3 reticulations (Fig S1) using a log likelihood ratio test. All the networks were visualized using Dendroscope3 (*89*). By comparing the fit of models with increasing number of reticulations, we assessed whether the inclusion of additional reticulations significantly improved the model fit. A low p-value (typically < 0.001) suggests that the added reticulation significantly enhances the fit in our data.

---

**Figure S1.**
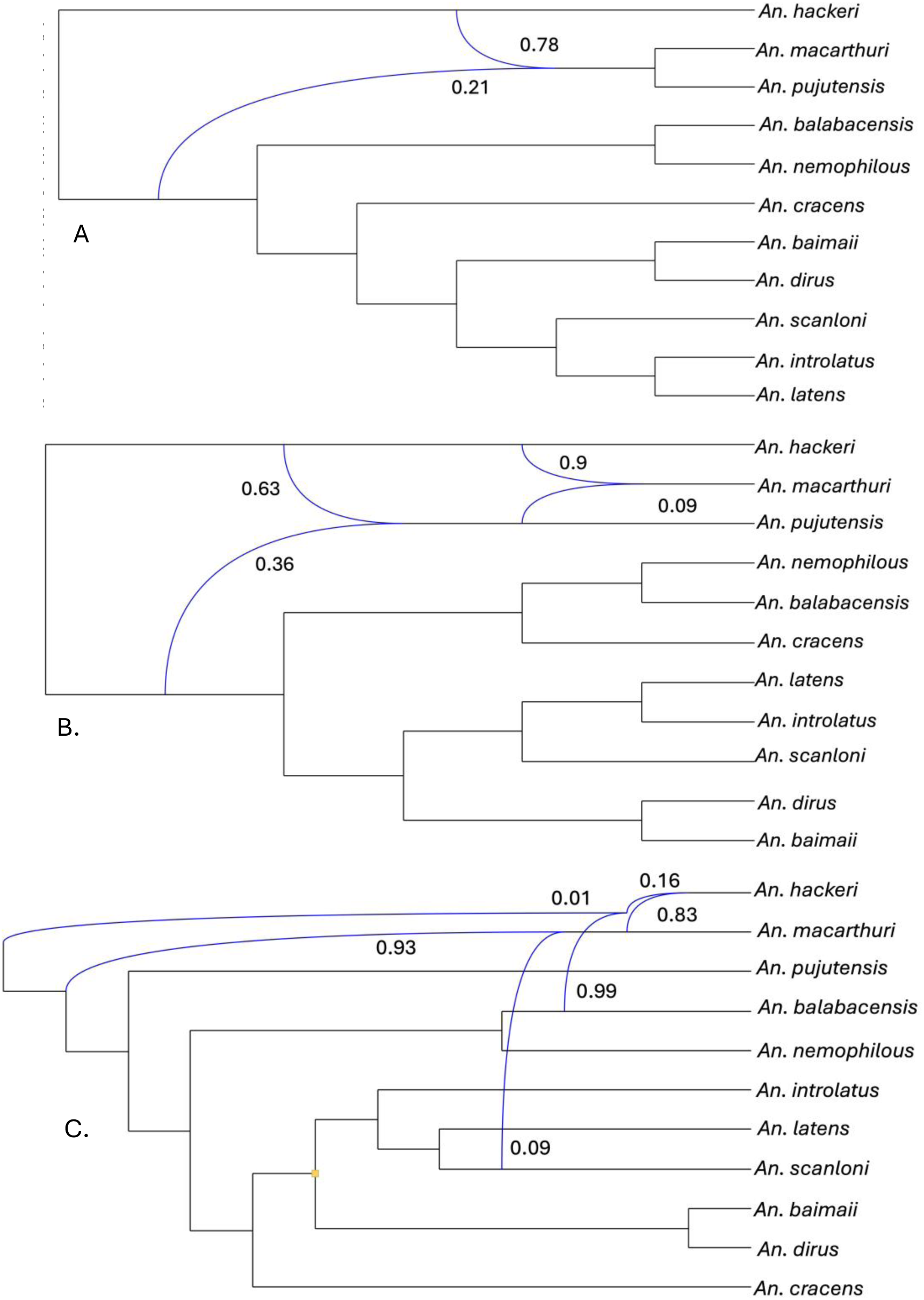
Phylogenetic networks inferred using maximum likelihood in PhyloNet. Optimal networks A, B and C were inferred when the number of reticulations was set to 1, 2 and 3, respectively. The log-likelihoods of the three networks are −1733.79 (**A**), −1727.66 (**B**), and −1715.53 (**C**) with p< 0.001. The backbone network is depicted in black solid lines. The reticulation edges are shown in blue lines. The numbers adjoining reticulation nodes are inheritance probabilities.

**Table S1.**
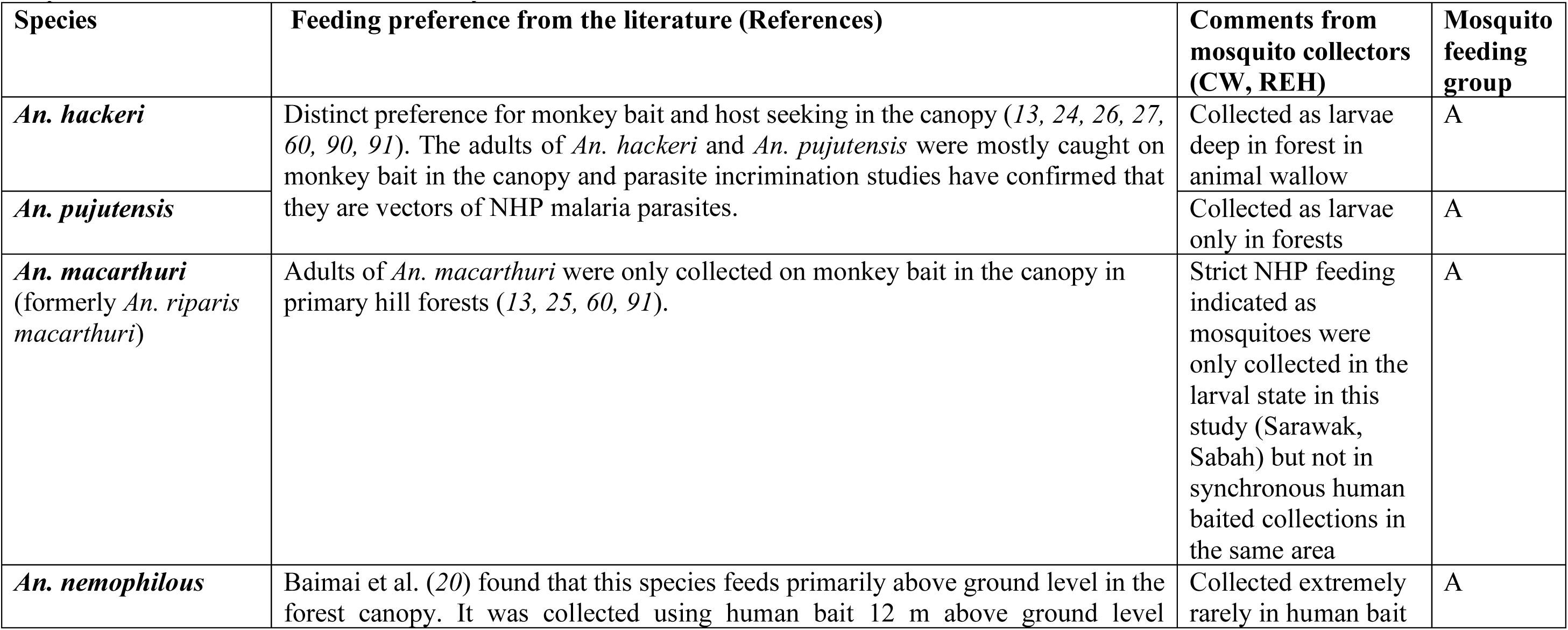
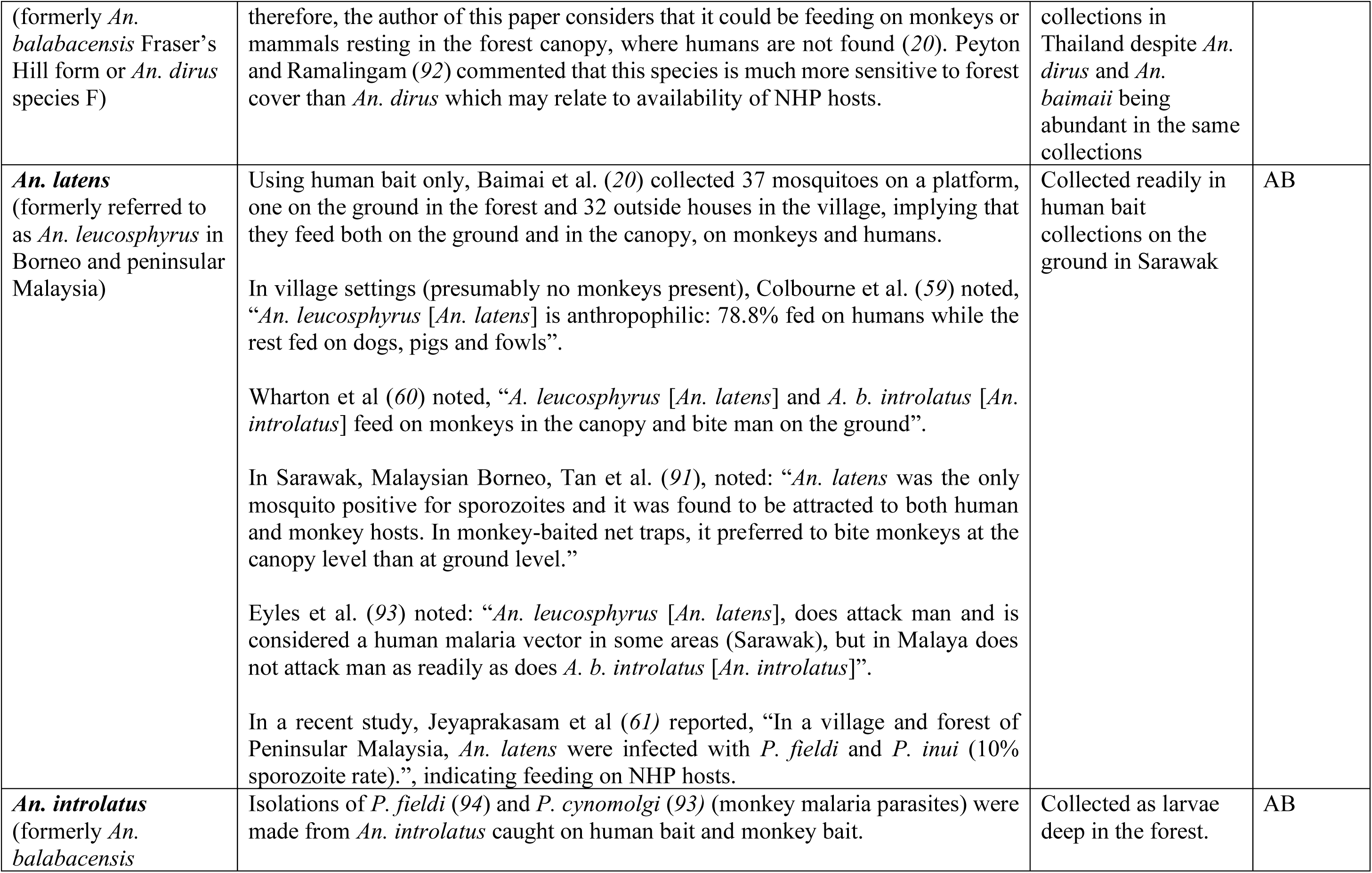
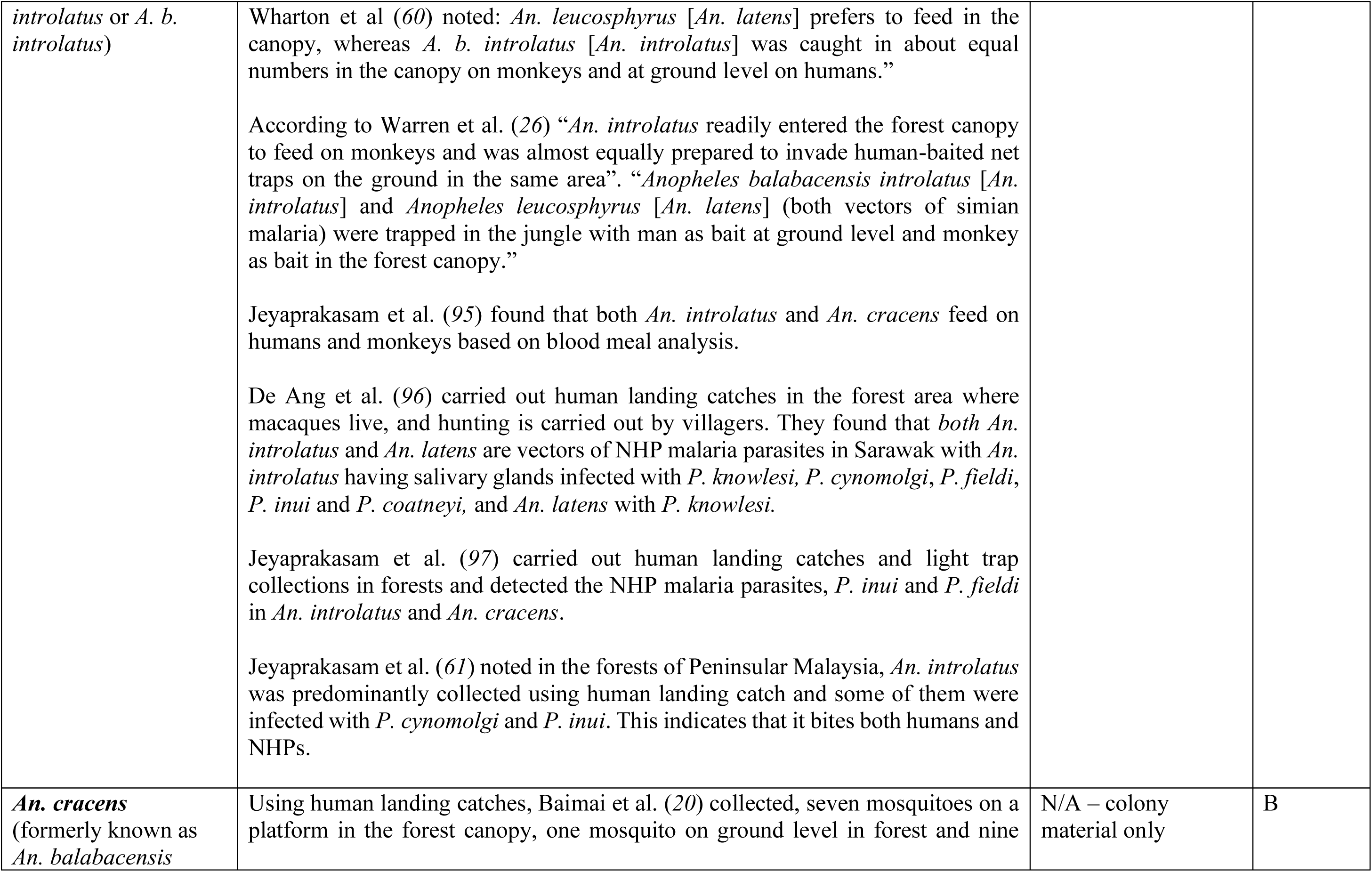
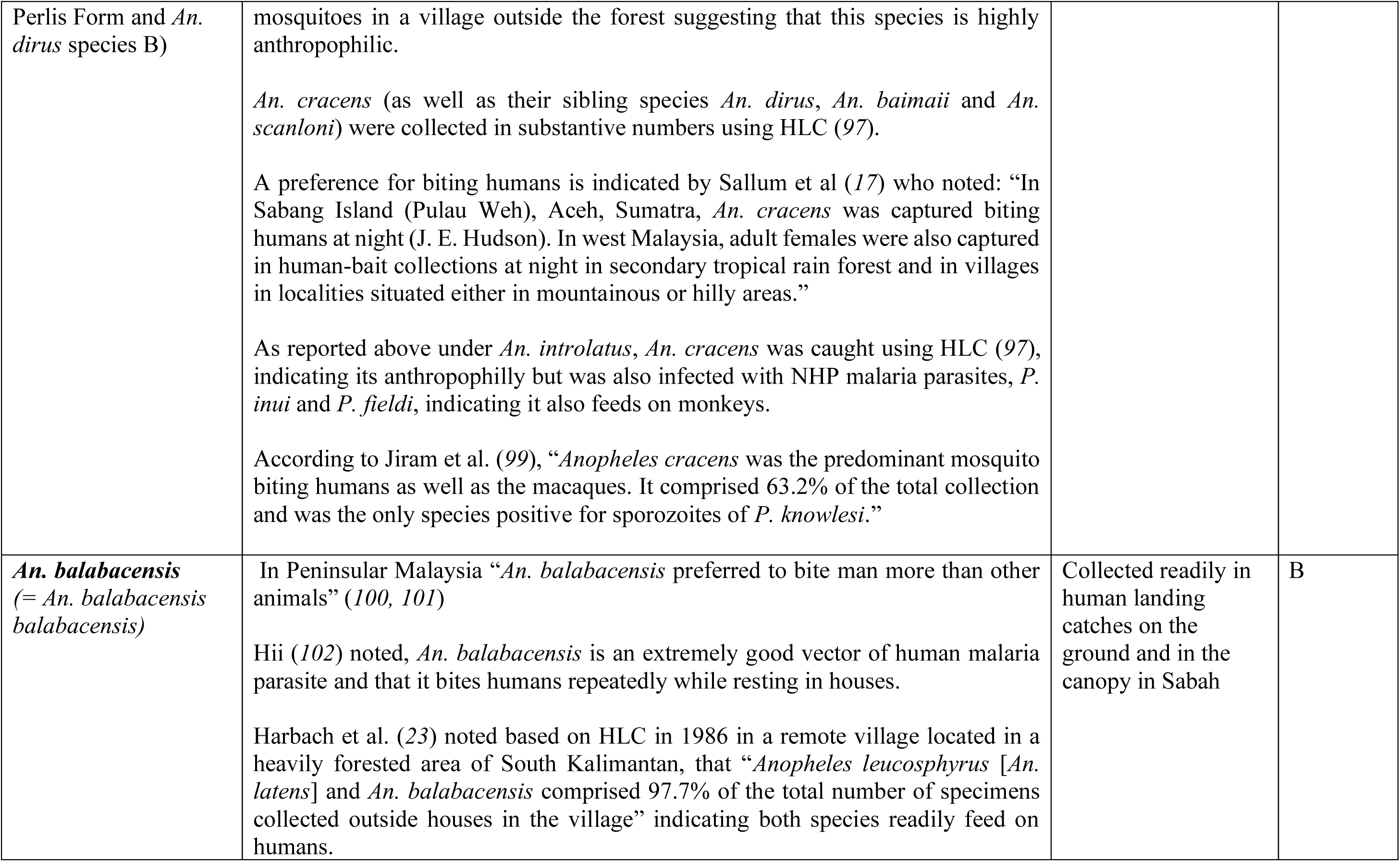
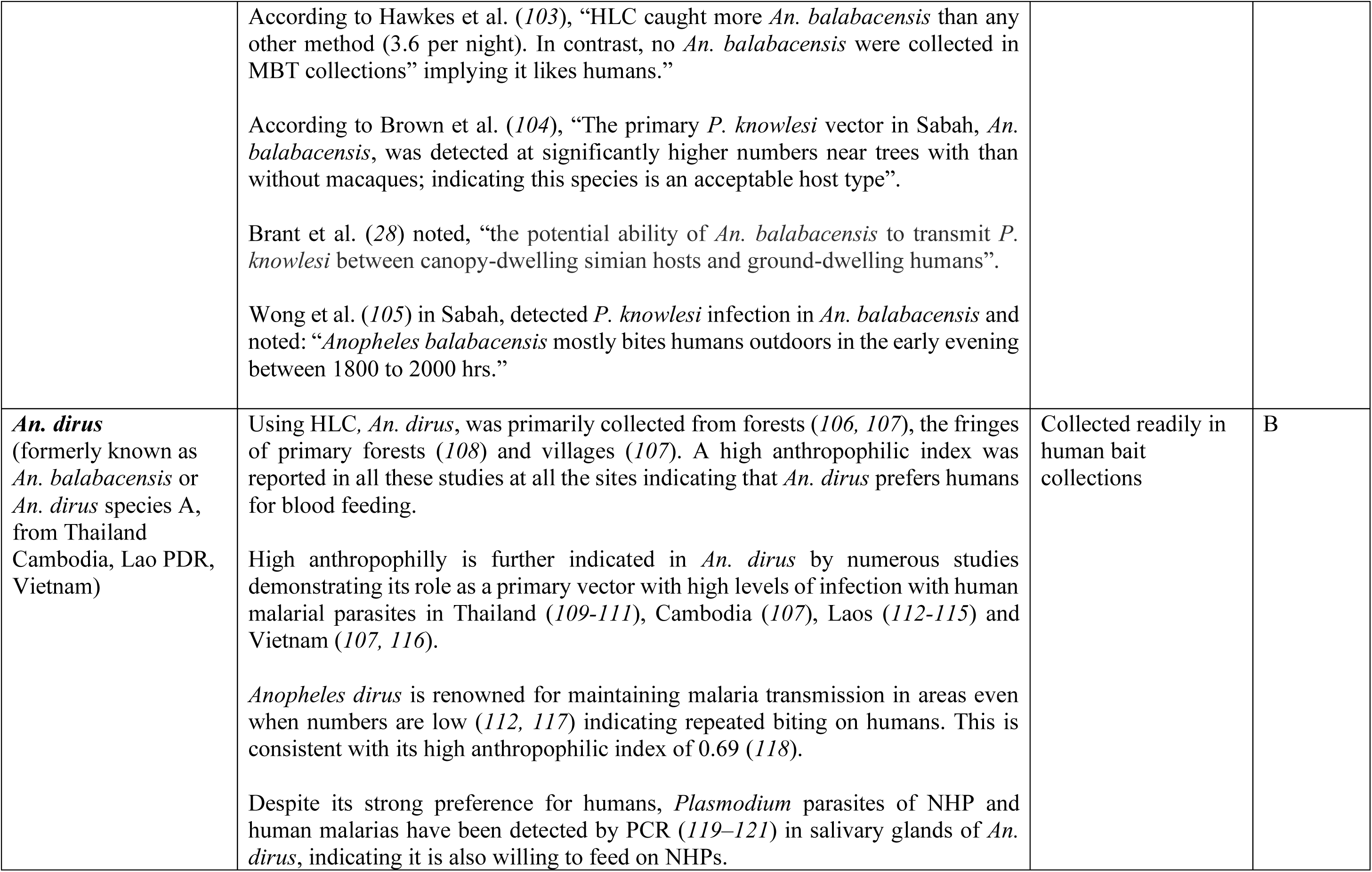
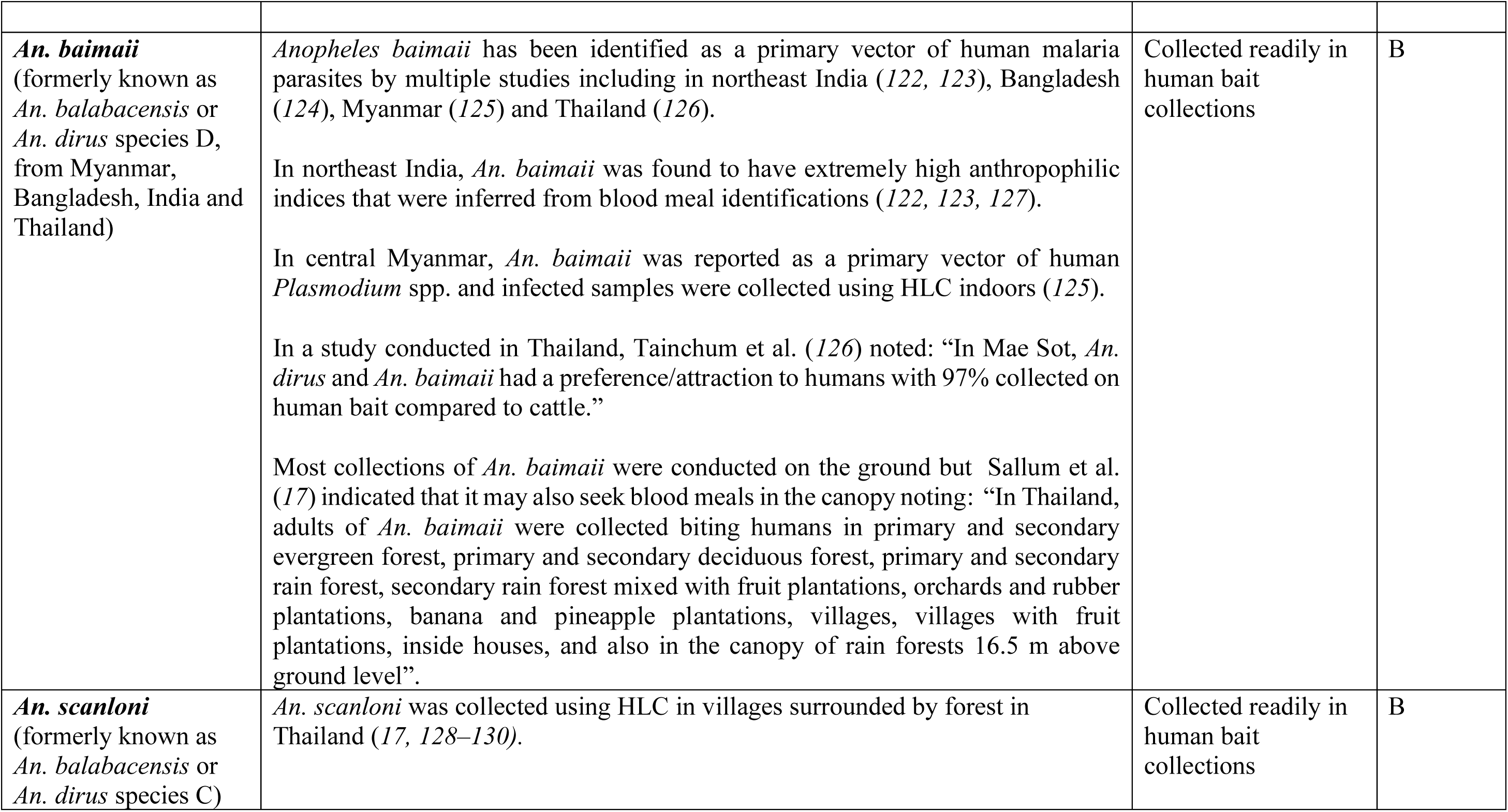
Evidence base for host feeding preferences of species of the Leucosphyrus Group used in this study. The table compiles information from studies that used monkey and human baits variously deployed in the canopy and on the ground to study the feeding behaviour/host preference of Leucosphyrus Group mosquitoes. It also includes comments relevant to host preference made by observations during field collections of mosquitoes used in this study. The last column indicates the feeding category into which species were placed based on this evidence as used in the text and analysis of host preference trait evolution in RASP v4. The three categories are: (A) strictly NHP feeders in the canopy; (B) highly anthropophilic behaviour at ground level and AB-host attraction for both NHP and humans. “Monkey feeding behaviour” is inferred here primarily from host choice experiments that were conducted using monkeys as bait (monkey bait traps [MBT]) however, as an extension we also imply host attraction (for blood feeding) to other NHPs in the forest of Southeast Asia such as gibbons and orangutans. Infection with monkey malarias (*P. knowlesi*, *P. cynomolgi*, *P. inui*, *P. coatneyi* and *P. fieldi*) is taken to indicate feeding on NHPs. HLC refers to human landing catches and where substantive numbers are caught indicates a preference for feeding on humans rather than NHPs in the canopy. Mosquito species names have changed over time as more species were discerned. While quotes from papers use the old name, the new name is indicated immediately afterwards in square brackets, where this can be reliably inferred.

**Table S2.**
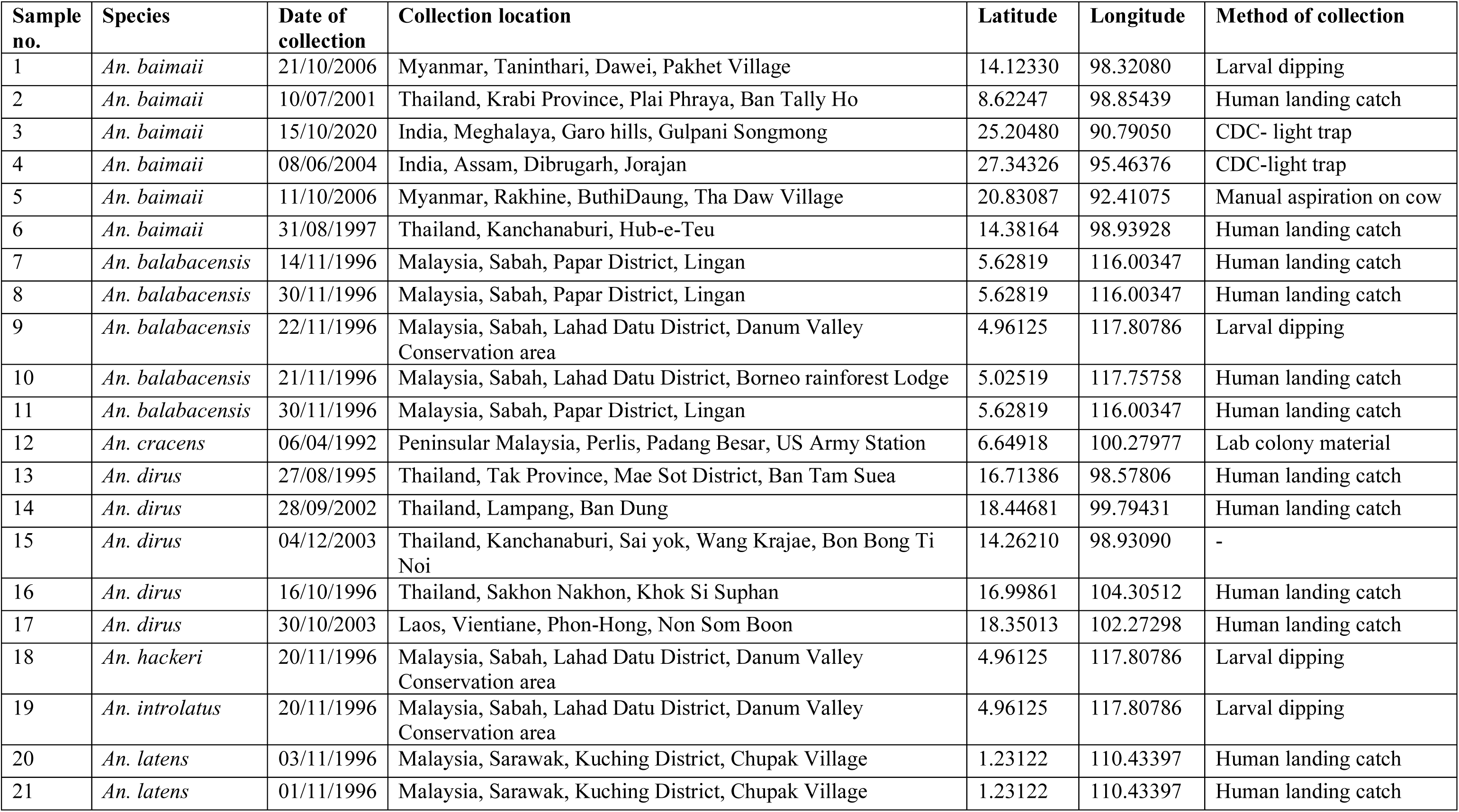
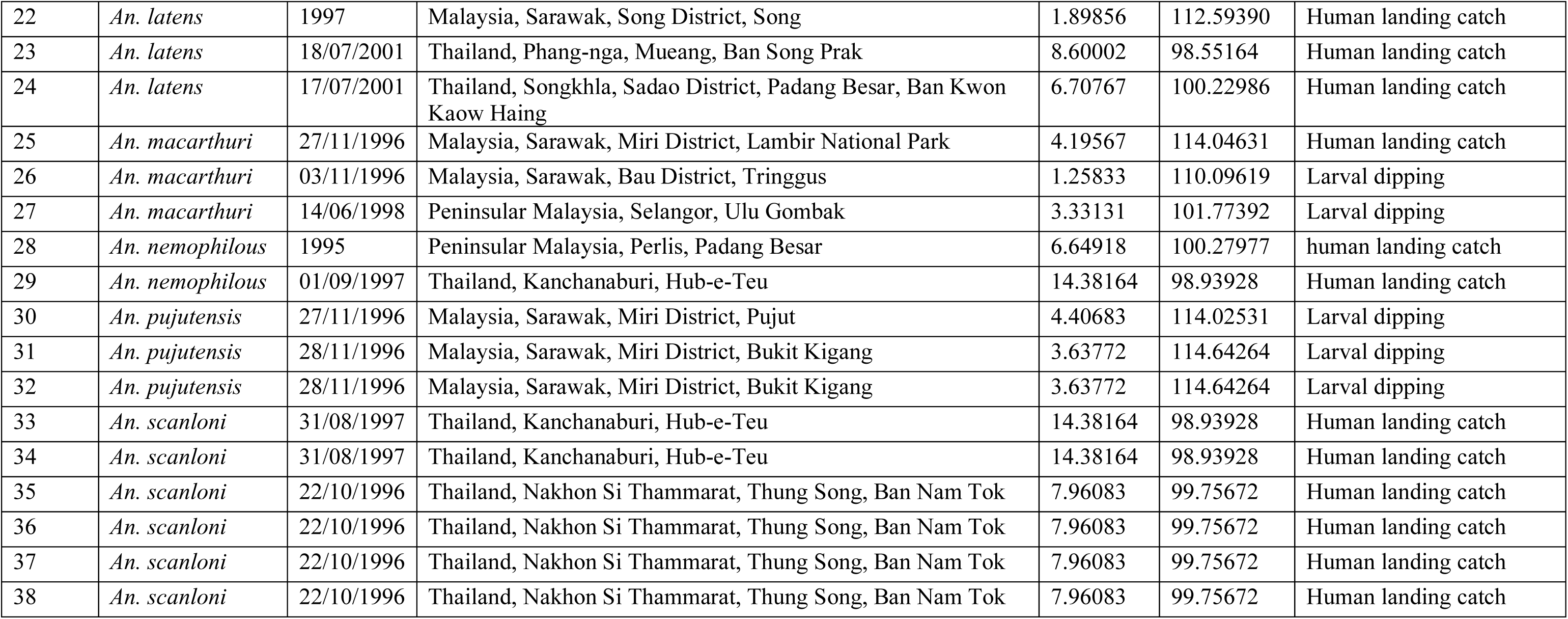
Details of mosquito samples, geographical locations and collection method used in the study.

**Table S3.** The current geographical ranges of members of the Leucosphyrus Group mosquitoes from the literature.

| Species | Geographical range |
| --- | --- |
| *An. macarthuri* | Peninsular Malaysia, Borneo, Thailand K-PL-ISK |
| *An. hackeri* | Borneo, Palawan, Peninsular Malaysia, Thailand K-PL-ISK |
| *An. pujutensis* | Sumatra, Peninsular Malaysia, Borneo, Thailand below K-PL |
| *An. balabacensis* | Palawan, (Sabah and Sarawak/Borneo), Java, Kalimantan, Brunei, |
| *An. latens* | Peninsular Malaysia, Borneo, Southern Thailand (below (ISK)) |
| *An. introlatus* | Peninsular Malaysia, Thailand below K-PL, Indonesia (in Sumatra), Borneo |
| *An. nemophilous* | Peninsular Malaysia, Thailand (K-PL-ISK), Northwestern Thailand, Thailand-central to north |
| *An. cracens* | Peninsular Malaysia, Thailand K-PL-ISK, Sumatra in Indonesia |
| *An. scanloni* | Northwestern Thailand, Thailand K-PL-ISK |
| *An. dirus* | Cambodia, Vietnam, Laos, Hainan-China, Thailand (central to north) |
| *An. baimaii* | Northeast India including Andaman and West Bengal, Myanmar, Bangladesh, Thailand (K-PL-ISK) + Northwestern Thailand |

## Notes

### Competing Interest Statement

The authors have declared no competing interest.

